# Phages use contingency loci as a bet-hedging strategy

**DOI:** 10.1101/2025.10.21.683753

**Authors:** Jasper B. Gomez, Jeffrey E. Barrick, Christopher M. Waters

**Author notes:** Corresponding Author: 5180 Biomedical and Physical Sciences 567 Wilson Road, East Lansing, MI 48824. current address-Department of Microbiology, Genetics, & Immunology, Michigan State University, East Lansing, Michigan, USA. Declarations of interest: none.

## Abstract

Bacteriophages are estimated to outnumber bacteria by ∼10-fold^1,2^. Here, we show that phage genomes contain contingency loci (CL), hypermutable DNA regions that promote reversible frameshift mutations through DNA polymerase slippage^3–6^. CL have been described in bacteria, archaea, and eukaryotes but have not previously been reported in phages. We demonstrate that CL in coliphage T2 and T4 generates genomic and phenotypic diversity in resulting progeny to evade host defense, a process known as bet-hedging. Whole genome sequencing of T2 and T4 show similar levels of CL-driven sequence variation in dozens of other putative CL. Additional sequencing of T6, T7, Secφ27, ICP1 and ICP2, alongside bioinformatics of the BASEL phage collection reveals that putative CL are widespread in phages and are encoded in every functional class of genes. Collectively, our study describes a new paradigm for understanding phage replication in which CL drive genetic diversification and population heterogeneity to rapidly evolve.

## Introduction

Bacteriophages (phages) encounter diverse bacterial hosts, and hundreds of bacterial phage defense systems, that enable hosts to survive infection^7–11^. To infect and replicate, phages must overcome such host defenses ^12^. We discovered that phage genomes rapidly adapt to such selective pressures through contingency loci (CL). A key feature of CL is the presence of DNA repeats that drive high mutation rates due to slipped strand mispairing by DNA polymerase during replication^4–6^. These hypermutable regions of DNA mediate high frequency, reversible, heritable, stochastic, genotypic switching^3,13^. CL have been widely studied in eukaryotes, bacteria, and archaea^13–15^; however, they have not been described in phages. Here, we demonstrate that phage CL drive diversity of progeny in a bet-hedging strategy to overcome bacterial defense. Furthermore, we find that CL are widespread in phage genomes, occurring in all functional gene classes, suggesting they mediate phage evolution to overcome other selective pressures in addition to host defenses.

### T2 mutants resistant to TgvAB rapidly evolve

TgvAB, a Type IV restriction system, is encoded in *Vibrio cholerae* ^16,17^. Heterologous expression of TgvAB in *Escherichia coli* protects it from T2, T4, and T6 phage infection by recognizing the glucosylated 5-hydroxymethylcytosine (5-hmC) on the phage DNA ^16,17^.

Addition of glucose to 5-hmC on phage DNA is encoded by the α-glucosyltransferase gene (*agt*) encoded in T-even phages^18^. T2 can escape TgvAB detection by acquiring null mutations in the *agt* gene ^16^. However, loss of glucosylation on 5-hmC results in an evolutionary trade-off as unglucosylated 5-hmC is recognized and restricted by the McrA/BC Type IV restriction enzymes ^16,19,20^ (Fig. 1A).

**Fig. 1.**
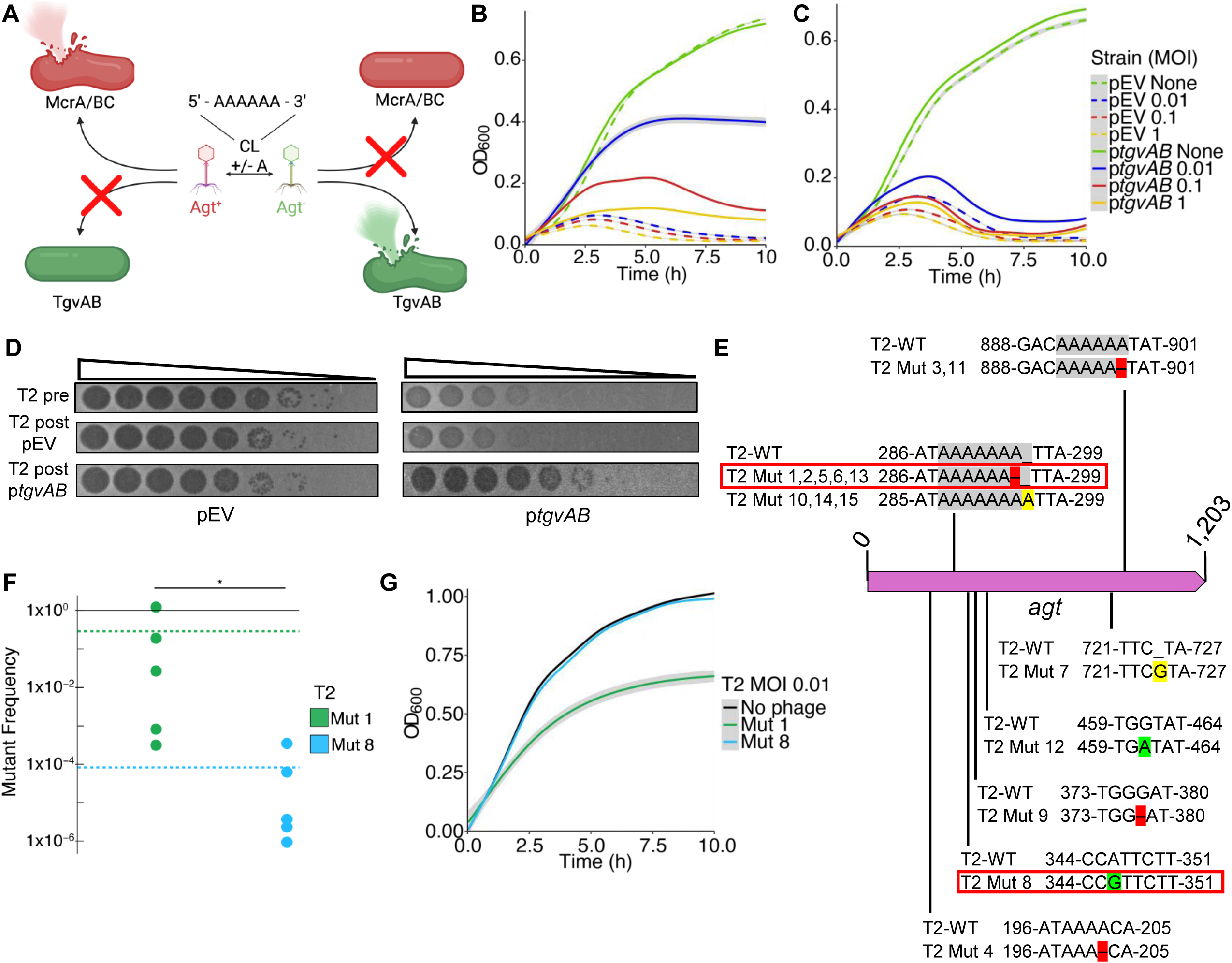
CL in T2 *agt* facilitates exploiting hosts with varying restriction systems. **A.** Schematic of WT *agt* or *agt* null mutant infection on either host with McrA/BC or TgvAB. **B.** *E. coli* with p*tgvAB* and pEV (empty vector) infected with WT T2 at varying multiplicities of infection (MOI’s) at time 0. The mean and standard error of 3 biological replicates each with 3 technical replicates are presented. **C.** T2 phage isolated from the cultures in Fig 1B was used to reinfect *E.coli* DH10B with p*tgvAB* and pEV at varying MOI’s at time 0 to determine if the phage had gained resistance. **D.** 10-fold serial dilution plaque assay of T2 pre-and post-infection on pEV and T2 pre-and post-infection on p*tgvAB* on host pEV and p*tgvAB.* The mean and standard error of 3 biological replicates each with 3 technical replicates are presented. **E.** Schematic of WT T2 *agt* gene. Nucleotide sequence of mutants and the surrounding region in *agt* compared to WT T2 sequence. Regions highlighted in red indicate a nucleotide deletion while yellow indicates a nucleotide insertion and green indicates a substitution in corresponding position. Mutants chosen for the reversion experiment are boxed in red. **F.** Mutant frequency of T2 Mut 1 (single strand repeat) and Mut 8 (point mutation). Dots represent one biological replicate, and dashed line represents the average of 5 biological replicates per T2 mutant. Wilcoxon test performed, p-value = 0.01 and is indicated by * **G.** MG1655 infected with T2 Mut 1 or T2 Mut 8 at an MOI of 0.001 at time 0. The mean and standard error of 5 biological replicates each with 3 technical replicates are presented.

During liquid culture infections, *E. coli* expressing TgvAB (p*tgvAB*) exhibited only transient protection against T2 during the initial stages of growth for all multiplicities of infection (MOI) (Fig. 1B). This dynamic differs from other restriction systems in which phage infection of hosts encoding these defenses does not significantly impact growth dynamics^10,11,21,22^. We hypothesized that the inhibition of growth by T2 in these cultures was due to rapid selection of TgvAB-resistant T2 mutants. To test this, we collected T2 phages after infecting *E. coli* (p*tgvAB*) and reinfected the same strain. These phages now exhibit complete killing of *E. coli* (p*tgvAB*), demonstrating rapid evolution of T2 resistance to TgvAB (Fig. 1C, D).

### T2 *agt* functions as a contingency locus

To identify which T2 mutations provide TgvAB resistance, we isolated spontaneous TgvAB-resistant plaques from 15 independently propagated T2 populations by plating WT T2 phages on *E. coli* (p*tgvAB*). Whole genome sequencing (WGS) revealed that phages from all 15 TgvAB-resistant plaques had mutations in *agt.* This result is consistent with our previous findings that null mutations in *agt* provide T2 resistance to TgvAB ^16^. Five mutants had unique single nucleotide polymorphisms (SNPs) in the *agt* gene while 10 mutants had frameshift mutations of +/-1 in two monomeric adenine (A) single sequence repeats (SSRs) (Fig. 1E). Due to the frequency of mutations in these *agt* SSRs and the rapid evolution of *agt* resistant T2 mutants, we hypothesized that these SSRs were CL in *agt*.

CL are repeat sequences in DNA that have an elevated rate of reversible mutations relative to the basal rate of DNA polymerase due to DNA polymerase slippage during replication^3,6^. To explore whether the *agt* SSRs exhibit an elevated reversible mutation rate, we quantified the *agt* reversion rates of T2 Mut 1 (deletion of an adenine in the *agt* SSR encoded from 288-296 bases into the *agt* gene) and T2 Mut 8 (an adenine to a guanine single nucleotide polymorphism (SNP)) (Fig. 1E). Equivalent numbers of phages from five independently harvested populations of each *agt* mutant were plated on *E. coli* strain MG1655 as it encodes the McrA/BC Type IV restriction system that recognizes and restricts *agt-* T2 phages (Fig. 1A), thus selecting for restoration of *agt* + phages. Our results from these five independent populations showed that T2 Mut 1 reacquired *agt+* at a frequency ∼1000-fold that of the T2 Mut 8 (Fig. 1F). Sequencing of phages from McrA/BC resistant plaques showed that both the SSR and SNP mutants produced true *agt* revertants. Moreover, T2 Mut 8 (SNP) was significantly more restricted by *E. coli* MG1655 in a liquid culture infection whereas T2 Mut 1 (SSR) more rapidly evolved resistance at all MOIs tested (Fig. 1G, Extended Fig. 1), illustrating the elevated reversion rate of the SSR allows rapid development of resistance to McrA/BC. These results indicate that the T2 SSR in *agt* at bases 288-296 is a CL.

### Putative CL are widespread in the T2 genome

Because CL have an elevated mutation rate compared to the basal DNA polymerase error rate^3,6^, we hypothesized that *agt* CL mutations could be detected in individual sequence reads generated from Illumina short read sequencing of phage genomes. This approach could quantify the generation of sequence variation independent of phenotypic selection. To test this hypothesis, five independent WT T2 populations were propagated on a non-selective host and sequenced to a depth of ∼2,000 fold. We identified every SSR (of any base) with a length ≥6 bases and analyzed individual sequencing reads to quantify the number of single-base insertions or deletions in SSRs. We found that four out of the five WT T2 populations had detectable mutations in the *agt* CL encoded at bases 288-296. The average mutation frequency for this *agt* CL across all five populations was 7 × 10^-4^ (total mutant sequences/total WT sequences, Fig. 2A), a mutation frequency that cannot be accounted for by the basal mutation rate of T2 DNA polymerase or errors in Illumina sequencing ^23–25^.

**Fig. 2.**
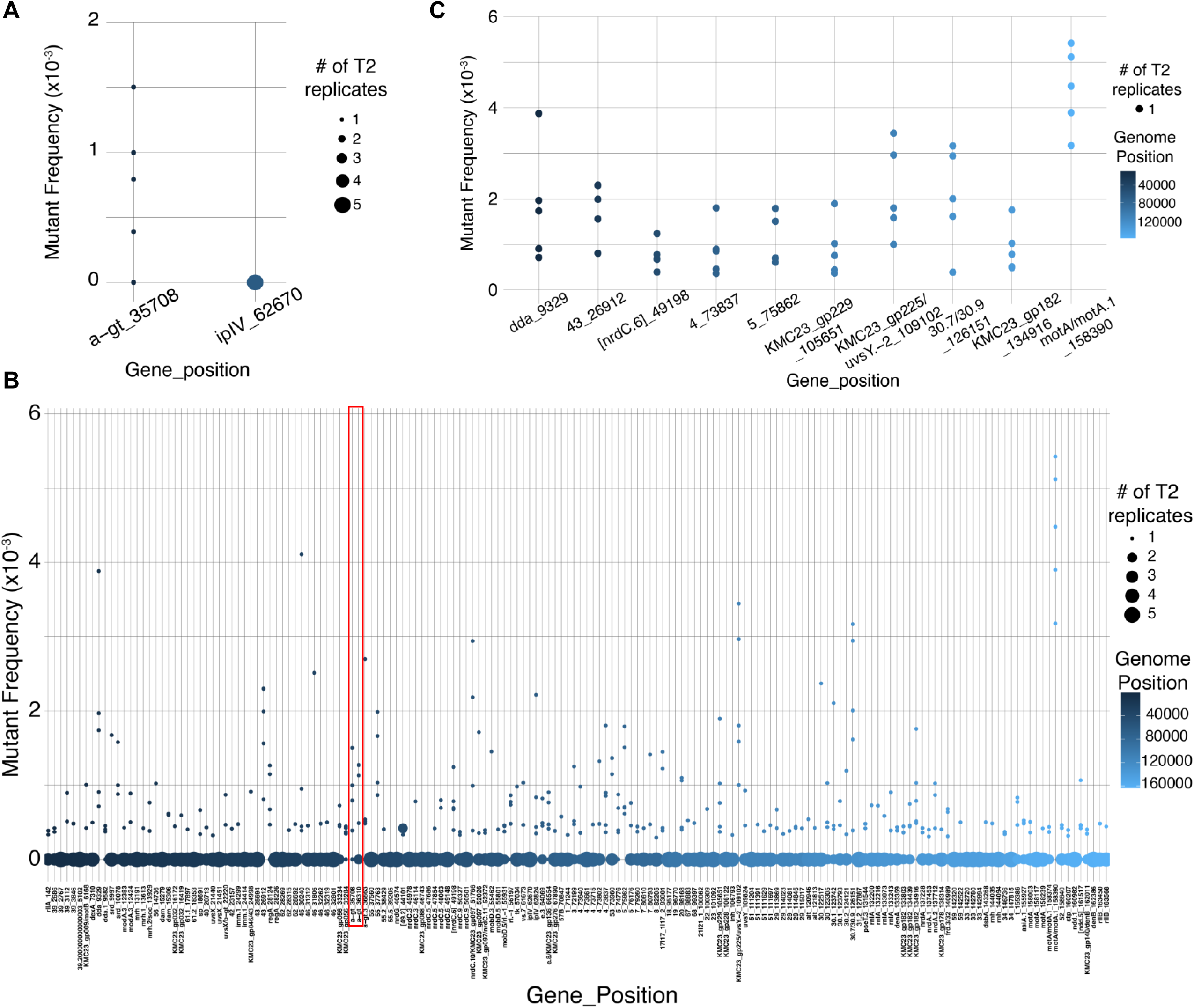
Variation in putative CL detected by Illumina Sequencing. 5 independent populations of WT T2 were sequenced. Sequencing reads with genomic regions with single sequence repeats (SSRs) ≥6 were analyzed and the number of reads that differed from WT sequence was calculated as Mutant Frequency. **A.** *agt* is shown with each dot representing an independent sequencing result. *ipIV* is shown with one enlarged dot representing all 5 independent sequencing results resulted in the same value of 0. Dot sizes correspond to number of independent sequencing with similar values. **B.** Percent mutants with all 5 independent sequencing values above 0. Slashes between gene names indicate the SSR is in an intergenic region between two ORFs. Color of dots represent position of repeat region in T2 genome. Start of genome is represented as dark blue while light blue indicates the end of genome. **C.** All SSRs in WT T2 plotted from the start of the genome (right) to end of the genome (left). The red box highlights the SSRs in *agt*.

Including the *agt* CL at bases 288-296, we identified 168 SSRs ≥6 bases spread across the genome of T2 (Fig. 2B and Supplemental Table 5) and found that 77.9% of SSRs had at least one population with detectable frameshift mutations from our whole genome sequencing analysis (Fig. 3A). Ten SSRs (5.95%) had mutations in all five populations that were sequenced (Fig. 2C, Supplemental Table 5). 22% of SSRs had no detectable mutations in any of the five mutant populations that were sequenced (Fig. 3A), and one example is shown in Fig. 2A. Mutations in individual sequencing reads could be generated during the Illumina library preparation or sequencing process. However, three pieces of evidence suggest this is not the explanation. First, the calculated average mutation rate determined from all 168 SSRs is 2.94 × 10^-4^, which is higher than the basal Illumina error rate^26^. Second, as mentioned 22% of SSRs had no mutations detected, suggesting that a repeat of ≥6 bases is not in itself sufficient to generate Illumina sequencing errors at this rate. And third, the mutation rates calculated from direct analysis of sequence reads were less than that determined by quantifying *agt*+ revertants (2.89 x 10^-1^, Fig. 1F) and a CL in *rIIA* described below (Fig. 4C), showing that biological quantification of CL mutation rates is on par with our sequencing read analysis. We therefore show that the 77.9% of SSRs with reliably detectable mutations are putative CL. The lack of observed mutations in 22% of SSRs could be due to inherent differences in mutation rates caused by *cis* acting sequences or by strong counter selection against mutations in these specific SSRs. Collectively, our results indicate that T2 encodes 131 SSRs that exhibit elevated mutation rates and are putative CL.

**Fig. 3.**
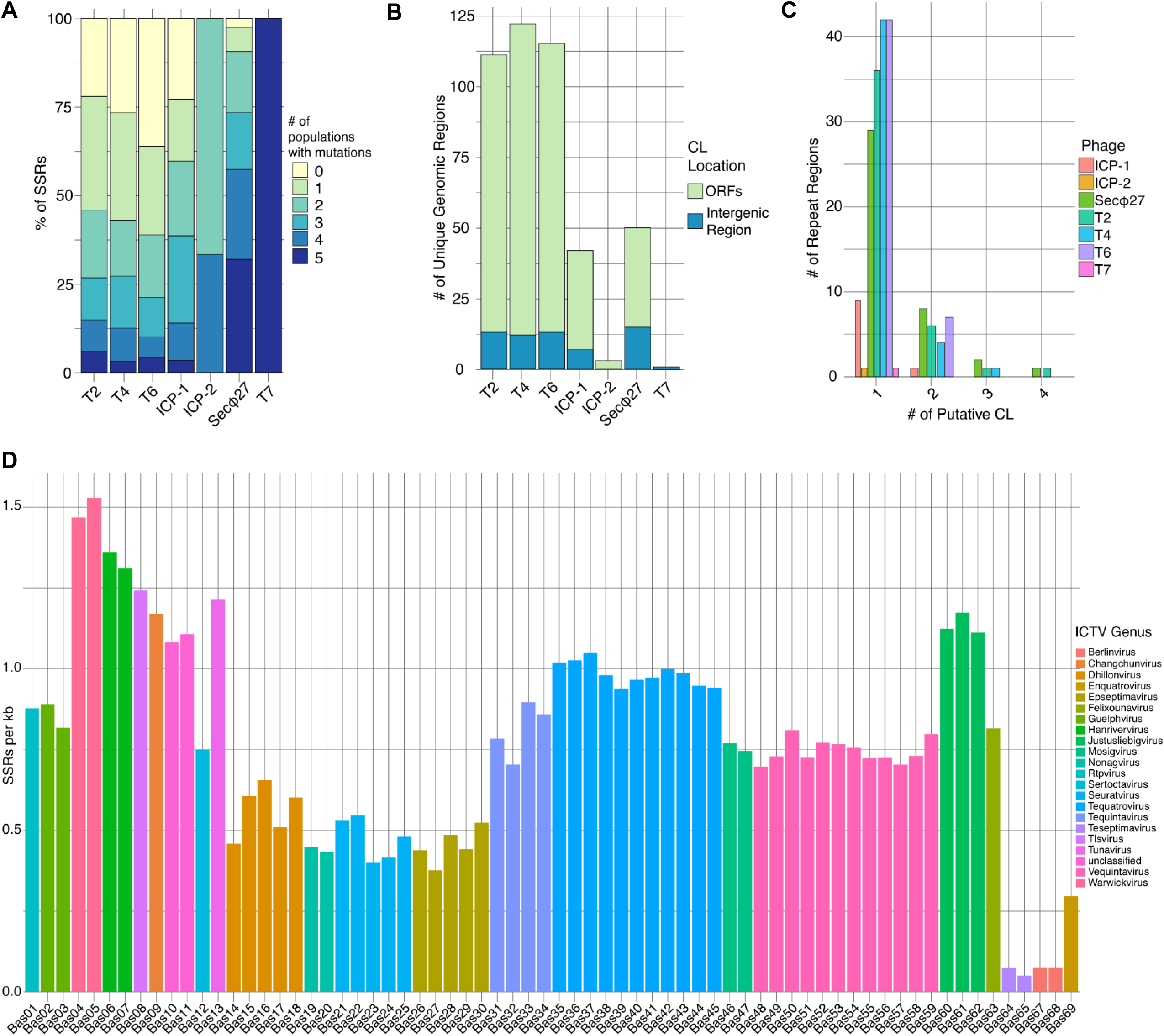
CL are widespread in other phages. **A.** Percentage of populations with mutations detected from 5 independent populations of T2, T4, T6, T7, Secφ27, ICP-1 and ICP-2 phages. **B.** Total number of unique genomic regions with putative CL in an open reading frame (ORF) or intergenic region for T2, T4, T6, T7, Secφ27, ICP-1 and ICP-2. **C.** Number of genomic regions (ORF/Intergenic region) with different numbers of putative CL for T2, T4, T6, T7, Secφ27, ICP-1 and ICP-2 genome. **D.** Number of SSRs per Kb identified in original BASEL phage collection.

**Fig. 4.**
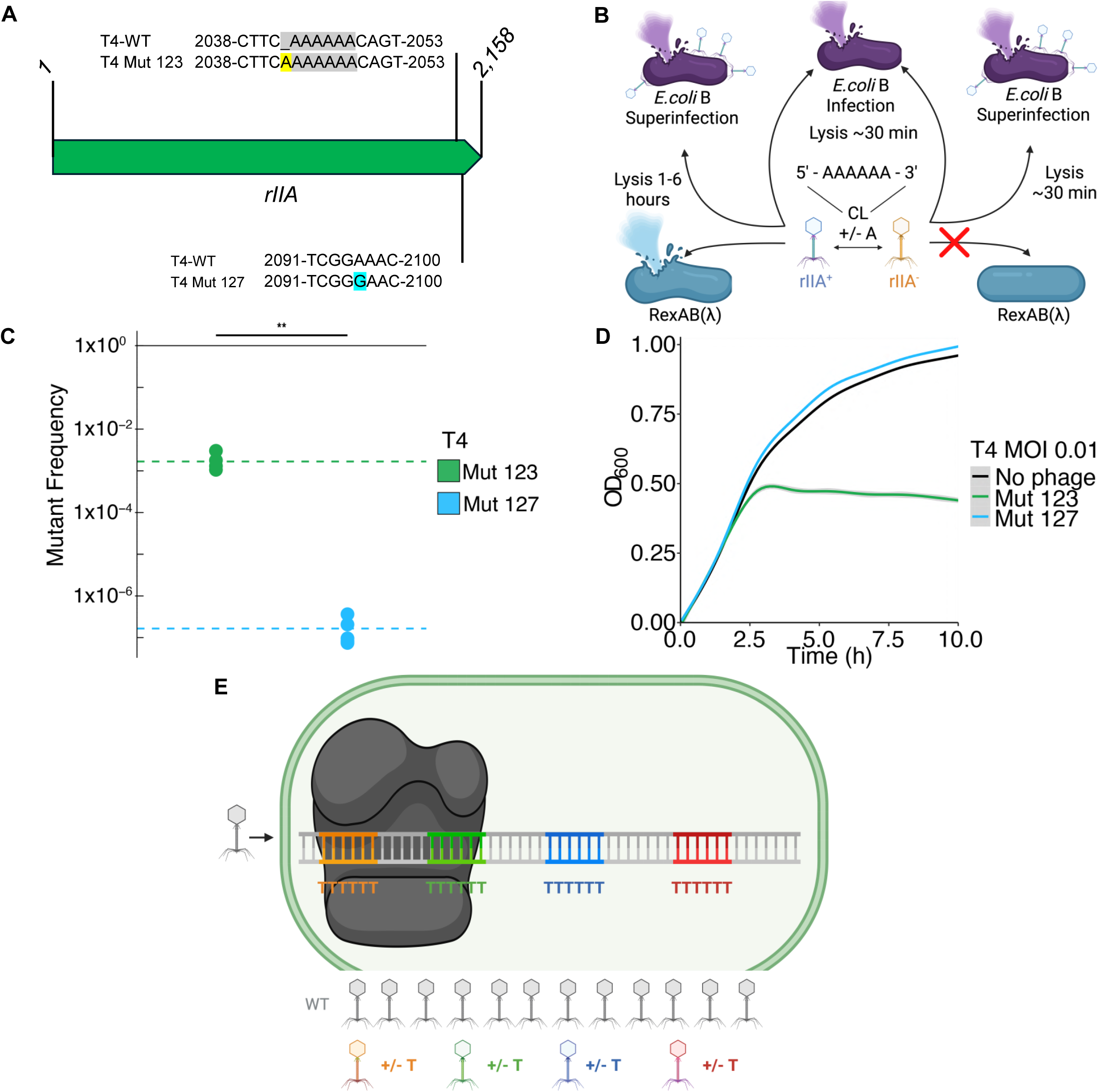
Evidence for CL reversion rate in T4 *rIIA*. **A.** Schematic of WT *rIIA* or *rIIA* null mutant infection on either *E.coli* B host with or without superinfection or host with RexAB (λ) **B.** Nucleotide sequence of mutants and the surrounding region in *rIIA* compared to WT T4 sequence. The region highlighted in yellow indicate a nucleotide insertion, and in blue indicate a substitution in corresponding position. **C.** Mutant frequency of T4 Mut 123 (CL) and Mut 127 (point mutation). Dots represent one biological replicate, and dashed line represents the average of 5 biological replicates per T2 mutant. Wilcoxon test performed, p-value = 0.007 and is indicated by ** **D.** MG1655(λ) infected with T4 Mut 123 or T4 Mut 127 at an MOI of 0.001 at time 0. The mean and standard error of 5 biological replicates each with 3 technical replicates are presented. **E.** Model for CL driven evolution in phages. When an individual phage replicates in its host, nucleotide repeat mutants arise through polymerase slippage at different CLs, resulting in a phage population with many distinct genomes.

### Diverse phages encode contingency loci

The T-even phages T2, T4, and T6 are morphologically, antigenically, and genetically similar^28–30^, and we hypothesized that the T4 and T6 phage genomes would also encode multiple putative CL like T2. To test this hypothesis, we propagated five independent populations each of T4 and T6, then sequenced their genomes and analyzed mutations in their SSRs as described above. We identified 191 SSRs in phage T4 and 188 SSRs in phage T6 (Supplemental Table 6, 7). Of these SSRs identified in T4 and T6, we observed 140 and 120 SSRs with detectable mutations, respectively, and we consider these putative CL. This result closely matched with our observations for phage T2, demonstrating that each of these phages encode over a hundred putative CL in their genome (Fig. 3A).

To explore whether genetically distinct phages encode putative CL, we sequenced five independent isolates of two additional coliphages, including the well-studied T7^31^ phage and the poorly studied SecΦ27^32^ phage, and the two vibriophages, ICP-1 and ICP-2^33–35^. We assessed the presence of SSRs by analyzing individual sequencing reads as described above. Our results identified SSRs for each phage; however, the number varied across phages with ICP-1 and SecΦ27 having 57 and 75 SSRs, respectively, while ICP-2 and T7 only had 2 and 1 SSRs, respectively (Supplemental Tables 8, 9, 10 and 11). Analogous to the T-even phages, the majority of SSRs had detectable sequencing reads with base insertions or deletions in at least one of the populations ranging from 77.2% in ICP1, 97.3% in SecΦ27, and 100% in ICP2 and T7 (Fig. 3A).

Most putative CL for all phages analyzed are present in open reading frames (ORFs) while a few are encoded in intergenic (IG) regions (Fig. 3B). Although this result is not surprising given that most of the phage genome is dedicated to encoding ORFs, it does suggest that many phage genes have the potential to be inactivated by frameshift mutations in CL. Additionally, we found that 100% of genomic regions in ICP-2 and T7, 90% in ICP-1, 89.4-81.2% in the T-even phages, and 72.5% in SecΦ27 had only one putative CL, but other genomic regions encoded 2-4 putative CLs, suggesting that such regions are mutational hot spots (Fig. 3C). We analyzed the predicted function of genomic regions with putative CL in each *E. coli* phage and found they occurred in all functional categories designated by PhageScope^36^ (Extended Fig. 2A, Supplemental Table 12).

### SSRs are widespread in phage genomes

To further ascertain the range of SSRs in phages, we analyzed the genome sequences of 106 dsDNA *E. coli* phages from the BASEL phage collection, which contains representatives of the major *E. coli* phage ICTV Genus^37^. Every phage ICTV Genus from the original BASEL library (Fig. 3D) and newly described expanded library (Extended Fig. 2B, Supplemental Table 13) encoded SSRs ≥6 bases and within each ICTV Genus, the density of SSRs per kilobase (kb) in the genome varied. Three phage ICTV Genus (*Teseptimavirus*, *Kayfunavirus*, and *Berlinvirus*) had the lowest average SSR/kb ranging from 0.02-0.07 while the phage ICTV Genus (Warwickvirus) had the highest SSR/kb at an average of 1.52. Consistent with our sequencing results, the *Tequatrovirus* genus, which includes the T-even phages, encoded the 8th highest average SSR/kb out of 32 genera, suggesting abundant SSRs, while the T7 family *Teseptimavirus* encoded the 31st lowest average SSR/kb, suggesting the loss of SSRs. We compared the average SSR/kb of each phage genus to the %GC in their genome but found no correlation, suggesting differences in SSR abundance are not due to base content (Extended Fig. 3, Supplemental Table 14).

To more broadly understand the role of SSRs in phage evolution, we examined what types of protein-coding genes contained SSRs ≥6 base pairs in a diverse set of 41 coliphage genomes from the BASEL collection (Extended Fig. 2B). The 41 phages selected exhibit <90% sequence identity to remove closely related phages (see Methods). We found that these SSRs were significantly underrepresented in infection proteins, such as tail fibers, and replication proteins, such as polymerases, with only about 0.77× as many SSRs in these genes as expected from randomizations tests (see Methods) (Extended Fig. 4, Bonferroni adjusted p < 0.001). This result is not surprising as these proteins are likely to be essential, and there may be selection against the evolution of hypermutable sequences that can cause frameshift mutations in their genes. On the other hand, we observed 2.03× as many SSRs as expected in the regulatory protein category, which includes phage repressors, and this overrepresentation was highly significant (Extended Fig. 4, adj p < 0.001). There were also excess SSRs in the hypothetical and unclassified protein category (Extended Fig. 4, adj p = 0.014), which includes the *agt* gene. These results suggest that SSRs have evolved in diverse phage genes to enable evolutionary bet-hedging to survive a variety of selective pressures.

### T4 *rIIA* encodes a contingency locus

To investigate the function of another putative CL, we examined the repeat of 6 A’s encoded in the *rIIA* gene in T2, T4, and T6 sequencing reads, which had detectable mutations in all three phages as determined by sequence analysis (Supplemental Tables 5, 6, and 7). The *rIIA* gene was key in elucidating fundamental aspects of molecular biology including determination of a triplet genetic code and demonstration of homologous recombination^38–42^, but a CL has never been described in this gene. To examine whether the putative CL in *rIIA* exhibits a higher reversion rate relative to other mutations, we engineered an addition of an adenine in the putative CL (T4 Mut 123) and a substitution of an adenine to a guanine 53 bp upstream (T4 Mut 127), regenerating previously identified *rIIA* null mutations ^40,43^ (Fig. 4A). During superinfection, when phages outnumber hosts, WT T4 activates lysis inhibition (LIN) to delay infection.

T4 *rIIA* mutants undergo rapid lysis, overcoming the delayed infection in *E.coli* B. However, the loss of *rIIA* results in an evolutionary tradeoff as these phages are now susceptible to RexAB, a phage defense system encoded by a lambda lysogen ^40,44–46^ (Fig. 4B). Thus, we quantified restoration of *rIIA+* in T4 Mut 123 and T4 Mut 127 from five independent populations by plating on *E. coli* K-12 (λ). T4 Mut123, which had a mutation in the CL, reverted at ∼10,000× higher frequency than the SNP mutation T4 Mut 127, confirming this SSR repeat in *rIIA* is a CL (Fig. 4C). Analogous to our results with *agt,* the *rIIA* CL provided increased fitness upon infection of K12(λ) as the RexAB defense system only exhibited transient protection against the *rIIA* CL null mutant, T4 Mut123, but complete protection against the *rIIA* SNP mutant, T4 Mut127, (Fig. 4D, Extended Fig. 5).

## Discussion

Our results indicate that during genome replication of dsDNA phages, DNA polymerase slippage at CL produces frameshift mutations at rates thousands of times higher than basal mutation rates (Fig. 4E). This process generates genetic diversity and provides a bet-hedging strategy that enables phages to survive diverse selective pressures, such as bacterial phage defense systems.

Results from analyzing whole-genome sequencing of various phages for ≥6 SSRs revealed putative CL in ORFs with a wide range of functions that include assembly, lysis, packaging, regulation, and replication. The prevalence of CLs in genes of many functions suggests this evolutionary strategy likely extends beyond evolving to overcome phage defense to other selective pressures such as divergent bacterial hosts or environments.

CL have never been ascribed to phages, however, *Fletchervirus* phages encode 7-11 guanine (polyG) tracts in receptor binding proteins, generating diversity that allows a sub-population to infect *Campylobacter* when its phase-variable receptor is not expressed^47^. These repeats were not designated CL nor was their mutation rate quantified. PolyG tracts were identified in a thymine methylase gene of bacteriophages BBP-1^48^ and in PB1-like phages in a baseplate gene^49^, but they were also not designated CL nor studied further.

The mutational variation that arises from phage CL may reflect a history of “Red Queen” coevolution between bacteria and phages, where each side must constantly evolve to counter evolution by the antagonist. By evolving reversible, elevated frameshift mutations in CL, phages can harness their short life cycle and high numbers of progeny to rapidly and reversibly sample genetic variation to enhance evolution. For example, using a simple probabilistic model F_mut_=1-(1-m)^r^, where m is the average mutation rate and r is the total number of repeats, we estimate using T2 as an example, that ∼1% of progeny produced during an infection will have a mutation in a CL. Thus, CL can generate vast genetic variation in phages that has not been appreciated nor explored.

## Methods

### Strains and growth conditions

Bacterial and phage strains used in this study are listed in Supplementary Table 1. Unless otherwise stated, cultures were grown in Luria broth (LB) at 37**°**C with shaking at 210 rpm or LB plates at 37**°**C and supplemented with the following antibiotics for plasmid maintenance as necessary: kanamycin (100 μg mL^-1^) and/or carbenicillin (100 μg mL^-1^). Coliphages were propagated by streaking on MMB agar (LB + 0.1mM MnCl_2_ + 5 mM CaCl_2_ + 5 mM MgCl_2_ + 0.5% agar) with *E. coli* DH10B at a 1:1,000 dilution of an overnight culture and incubated overnight. Plates were then flooded with 10 mL of phage buffer (0.1 mM Tris HCl 7.5 pH + 10mM MgSO_4_ + 0.4% NaCl_2_ + dH_2_O) and incubated statically at 4**°**C for 24 hours. Phage buffer was then filtered through a 0.22 μM filter. Vibriophages were propagated exactly like coliphages phages except the host is *V. cholerae* E7946.

### Plasmid construction

Plasmids are listed in Supplementary Table 2 and primers in Supplementary Table 3. All PCR products were amplified using Q5-High Fidelity DNA polymerase according to the manufacturer’s instructions (New England Biolabs). All plasmid constructs were generated using fast cloning as described in Gomez et al. 2024^16^. All plasmids and PCR products were confirmed by sequencing (Plasmidsaurus). Plasmid sequencing results were mapped to T4 reference genome (NC_000866.4) using Geneious Prime 2023.0.2.

To construct pJBG122, *rIIA* was amplified from wild-type T4 Carolina with overlapping ends to pUC19 using oJBG206 and oJBG207. pUC19 was linearized using HF_pUC19-MCS_F and HF_pUC19-MCS_R. To construct pJBG123 and pJBG127, one region of pJBG122 was amplified with site-directed mutagenesis primer oJBG208 (pJBG123) or oJBG216 (pJBG217) with oMJF006 and another region was amplified with site-directed mutagenesis primer oJBG209 (pJBG123) or oJBG217 (pJBG127) with oMJF005. Both PCR products were used for fast cloning to create pJBG123 or pJBG127, respectively. Inserts were verified by PCR using primers oJBG191 and oJBG192.

### T4 *rIIA* site directed mutagenesis

To generate T4 Mut 123 and Mut 127, *E. coli* strain DH10B containing pJBG123 and pJBG127 respectively, were grown overnight in LB at 37**°**C. Overnight cultures were mixed 1:1,000 with 20 mL of MMB agar + carbenicillin and poured into 150 x 15mm Petri dish. Plates were left to solidify for 30 minutes at room temperature. After 30 minutes, 150 μl of wild-type T4 was spread across the plates. Plates were dried and placed at 37**°**C overnight. Plates were covered with 10 mL of phage buffer and placed at 4**°**C overnight. Phage buffer was collected and filtered using a 0.22 μM filter. Lysates were then taken and spread on plates made with an overnight culture of *E.coli* B at 1:1,000 and mixed with 20 mL of MMB agar and poured into a 150 x 15mm Petri dish and incubated at 37**°**C overnight. Plates were covered with 10 mL of phage buffer and placed at 4**°**C overnight. Phage buffer was collected and filtered using a 0.22 μM filter. Lysate was then streaked on plates made with an overnight of *E. coli* B at 1:1,000 and mixed with 10 mL of MMB agar to obtain isolated plaques. Plaques were then picked based on the large plaque phenotype on *E. coli* B. T4 Mut 123 and Mut 127 were verified by plating on MG1655(λ) and with PCR using primers oJBG191 and oJBG192.

### Phage infection and reinfection in liquid culture

Overnight cultures of DH10B p*tgvAB* and pEV (pJBG078 and pMJF103) grown in LB + kanamycin were diluted to an OD_600_ of 0.01 into 10 mL flasks containing LB + kanamycin. Cultures were grown for ∼2 hours to an OD_600_ 0.1. The cultures were distributed into a 96 well plate and wild-type T2 was added at MOIs of 0.1, 0.01, and 0.001. Infection was monitored by measuring OD_600_ every 2.5 minutes for 10 hours at 37**°**C using a BioTek 800 TS (Agilent) plate reader with continuous, linear shaking. After 10 hours plates were placed in 4**°**C. Cultures from the wells were then collected and transferred to a AcroPrep Advance 96-well filter plate for aqueous filtration with a 0.2 μM filter. The 96-well filter plate was placed on top of a another 96 well plate to collect phage lysate. Plates were centrifuged at 4,000rpm at 4**°**C. Phage concentration was calculated from infection on DH10B. Overnight cultures of DH10B p*tgvAB* and pEV grown in LB + kanamycin were diluted to an OD_600_ of 0.01 into 10 mL flasks containing LB + kanamycin. Cultures were grown for ∼2 hours to an OD_600_ 0.1. Once OD_600_ = 0.1, cultures were distributed into a 96 well plate. Using phage lysate from the initial *E. coli* p*tgvAB* infection, MOIs initially at 0.1, 0.01 and 0.001 were added again at the exact MOI into 96 well plate and measured with the same parameters as initial infection. Experiments were replicated 3 times independently and representative images are shown.

### Phage infection in liquid culture

Overnight cultures of MG1655 or K-12 (λ) were diluted to an OD_600_ of 0.01 into 10 mL flasks containing LB. Cultures were grown for ∼1.5 hours to OD_600_ 0.1. Once OD_600_ = 0.1, cultures were distributed into a 96 well plate and T2 Mut 1 and Mut 8 were added to MG1655 or T4 Mut 123 and Mut 127 were added to K-12 (λ) at MOIs of 0.1, 0.01, and 0.001. Infection was monitored as described above. Experiments were replicated 3 times independently and representative images are shown.

### Efficiency of plaquing (EOP) assays

Assays were performed as described in Gomez et al^16^. Briefly, overnight cultures of DH10B p*tgvAB* and pEV were mixed 1:1,000 with 20 mL MMB + kanamycin and poured into 150 x 15mm Petri dishes. Wild-type, post, and pre T2 stocks were diluted 1:10 up to a 10^-12^ dilution. 5 μl of each dilution was spotted on DH10B p*tgvAB* and pEV plates. Plates were dried before placing at 37**°**C overnight. Experiments were replicated 3 times independently and representative images are shown.

### Mutant frequency assay

T2 Mut 1 and T2 Mut 8 were struck out on overnight cultures of p*tgvAB* mixed 1:1,000 with 10 mL of MMB agar + kanamycin and poured into 100 x 15mm Petri dish. 5 isolated plaques were picked and resuspended in 150 μl of distilled H_2_O (dH_2_O). Overnight cultures of DH10B were mixed 1:1,000 with 20 mL of MMB agar and poured into 150 x 15mm Petri dish. Plates were left to solidify for 30 minutes at room temperature. After 30 minutes, 150 μl of each isolated plaque was spread, dried and placed at 37**°**C. Plates were covered with 10 mL of phage buffer and placed at 4**°**C overnight. Phage buffer was collected and filtered using a 0.22 μM filter.

T4 Mut 123 and T4 Mut 127 were struck out on overnight cultures *of E. coli* B mixed 1:1,000 with 10 mL of MMB agar + kanamycin and poured into 100 x 15mm Petri dish. 5 isolated plaques were picked and resuspended in 150 μl of distilled H_2_O (dH_2_O). Overnight cultures of MG1655 were mixed 1:1,000 with 20 mL of MMB agar and poured into 150 x 15mm Petri dish. Plates were left to solidify for 30 minutes at room temperature. After 30 minutes, 150 μl of each isolated plaque was spread, dried and placed at 37**°**C. Plates were covered with 10 mL of phage buffer and placed at 4**°**C overnight. Phage buffer was collected and filtered using a 0.22 μM filter. EOP of all isolated phage populations for T2 Mut 1 and Mut 8 were determined on MG1655 and DH10B. EOP of T4 Mut 123 and Mut 127 were determined on MG1655 and K-12 (λ).

Using EOPs we performed plaque-forming unit quantification assays as described in Gomez et al^16^ for all independent phage lysates. The following equation was used to calculate mutant frequency, m = ((D1/C1)*(C2/D2)) where D1 is the dilution factor from non-selective host lysate, C1 are PFU counts on non-selective host, D2 is the dilution factor from selective host lysate and C2 are PFU counts on selective host and m is the mutation rate. Experiments for each individual population were replicated 3 times independently and representative images are shown.

### Illumina sequencing

*E. coli* phage DNA extraction was done as described in Gomez et al. 2024^16^. Vibriophage extractions were done as described in Minmin et al. 2017^50^. High-titer phage lysates (>10^6^ PFU μl^-1^) were used for extraction. DNA extracted samples were sent to Seqcoast for short read whole genome Illumina sequencing. Short read whole genome Illumina sequencing was performed using their small 200 Mbp/1.3 million reads service with 150 bp paired end reads. Sequencing was performed using either a standard SBS flow cell or a XLEAP flow cell on a NextSeq 2000.T2 *agt* null mutations information is provided in Supplemental Table 4.

### Breseq analysis

Illumina sequencing results were analyzed using *breseq*^51^ with runs automated using *brefito*^52^. Sequences were aligned with their corresponding reference genome T2 (NC_054931.1), T4 (NC_000866.4), T6 (NC_05407.1), T7 (NC_001604.1), Secφ27 (NC_047938.1), ICP-1 (MH310933), and ICP-2 (NC_015158). Sequence information is provided in Supplemental Tables 5, 6, 7, 8, 9, 10 and 11. Genomic regions were pulled based on SSRs ≥6 using the tabulate-CL utility command of *breseq* in strict mode, which requires an exact match in a read to the five bases on either side of the SSR to count a read. Mutant frequency detected in the population was calculated by N/T where N = total number of mutant reads and T = the total number of reads. All analyses were done for 5 independent populations of each phage strain.

### BASEL phage CL analysis

Phage genomes from the BASEL collection were downloaded using GenBank accessions provided in Humolli et al 2025^37^. Putative CL in each genome were identified by using a Python script to scan for ≥6 base pair homopolymer SSRs. Gene contexts (ORF/intergenic region) of SSRs were classified based on annotations loaded from GenBank files using BioPython^53^.

GenBank accession numbers and SSR information are provided in Supplemental Table 13.

### Association of putative CL with phage gene categories

Coliphage genomes from the BASEL collection were analyzed^37^. First, we de-duplicated this set to 41 genomes that all had a pairwise average nucleotide identity (ANI) <90% as calculated using FastANI (v1.34)^54^. GenBank accession numbers and information about these sequences are provided in Supplemental Table 15. We transferred annotations of phage protein categories from the GenBank portion of PhageScope^36^ to proteins in these genomes via BLASTP (v2.15.0 +) searches^55^, taking top matches and requiring 70% query covered and an E-value 1E-6 to assign a category to a protein. Because there were few proteins in the “immune” and “tRNA_related” categories, we combined those with “hypothetical” and “unsorted” proteins into one “Other or unknown” category. Finally, we assessed the statistical significance of depletion or enrichment of SSRs (homopolymer SSRs ≥6 base pairs or longer) in the sequence of proteins in specific functional categories using randomization tests. For each randomization, we selected a new number of SSRs from a Poisson distribution with the actual number observed in the phage genomes as the mean, then shuffled SSRs among proteins in these phages, weighting by gene length. We repeated this procedure 20,000 times for each of eight categories. Two-tailed p-values for rejecting the hypothesis that the observed number of SSRs in a protein category would be observed by chance were calculated as twice the number of trails with the same or a more extreme number of SSRs compared to the actual genomes and Bonferroni adjusted by multiplying by the number of categories tested before assessing statistical significance.

## Acknowledgments

We thank Elizabeth N. Ottosen and Micah Ferrell for sharing primers used to construct plasmids, Chris Adami for consultation regarding calculating the percentage of phage population containing a CL mutation, and Bonnie Bassler and Richard Lenski for advice and comments on this paper. This research was supported and funded by NIH grants GM139537 and AI158433 to C.M.W, GM088344 to J.E.B., and F31AI186463 to J.B.G., and NSF grants DEB-1813069 and DEB-1951307 to J.E.B.

## Author contributions

J.B.G. performed the experiments, J.B.G. and J.E.B. analyzed the data and performed computations analysis, J.B.G. and C.M.W. wrote the manuscript and J.E.B. edited the manuscript.

## Supplemental Tables List

Table S1 – Strains used in this study

Table S2 –Plasmids used in this study

Table S3 – Primers used in this study

Table S4 – Illumina sequencing results of 15 independent T2 mutants passaged in presence of ptgvAB

Table S5 – Wildtype T2 SSR ≥6 Illumina whole genome sequencing reads

Table S6 – Wildtype T4 SSR ≥6 Illumina whole genome sequencing reads

Table S7 – Wildtype T6 SSR ≥6 Illumina whole genome sequencing reads

Table S8 – Wildtype ICP2 SSR ≥6 Illumina whole genome sequencing reads

Table S9 – Wildtype secΦ27 SSR ≥6 Illumina whole genome sequencing reads

Table S10 – Wildtype ICP1 SSR ≥6 Illumina whole genome sequencing reads

Table S11 – Wildtype T7 SSR ≥6 Illumina whole genome sequencing reads

Table S12 – Predicted gene functions of genomic regions with putative CL in *E.coli* phages

Table S13 – Total number of SSRs and SSRs per kb in BASEL phage library

Table S14 – BASEL phage ICTV Genus average SSRs and average GC content

Table S15 – 41 BASEL phage with ANI <90% sequence identity

---

**Extended Fig. 1.**
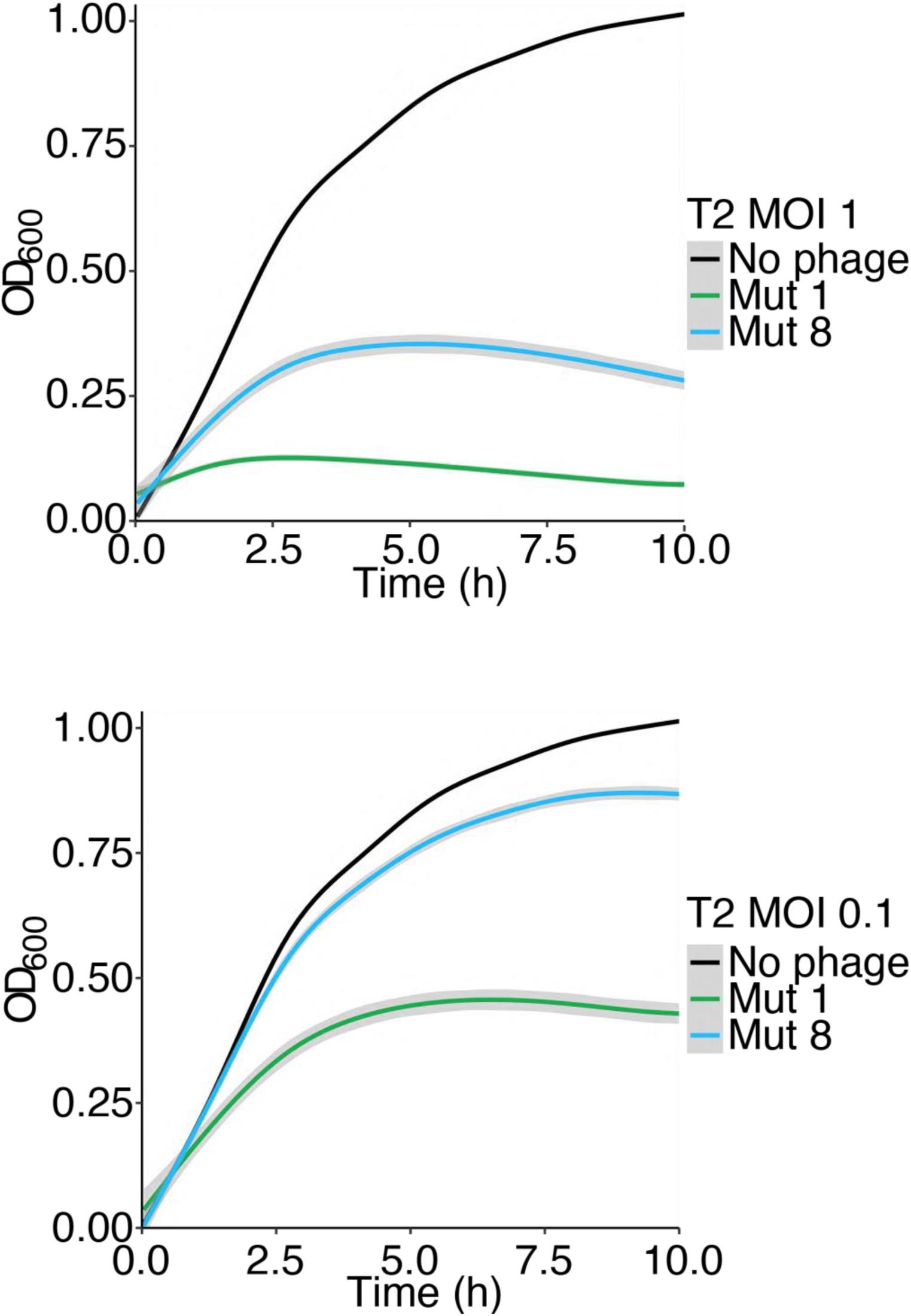
Contingency loci in T2 *agt* protects against varying restriction systems. MG1655 infected with T2 Mut 1 or T2 Mut 8 at an MOI of 1 and 0.1 at time 0. The mean and standard error of 5 biological replicates each with 3 technical replicates are presented.

**Extended Fig. 2.**
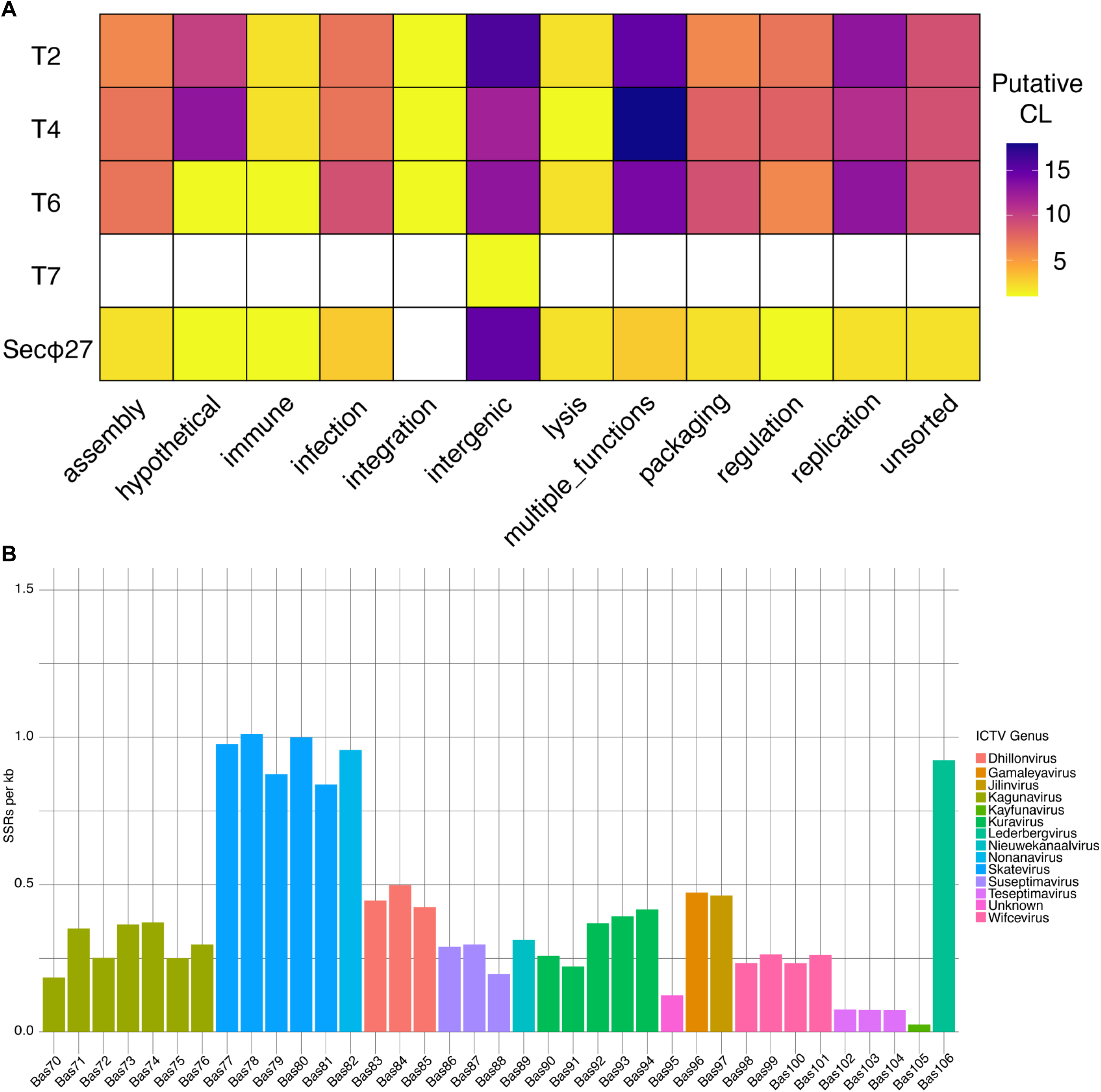
Simple sequence repeats ≥6 bases identified in other phages. **A.** Functional regions with putative contingency loci for T2, T4, T6, T7 and Secφ27 **B.** Number of SSRs per kb identified in extended BASEL phage collection.

**Extended Fig. 3.**
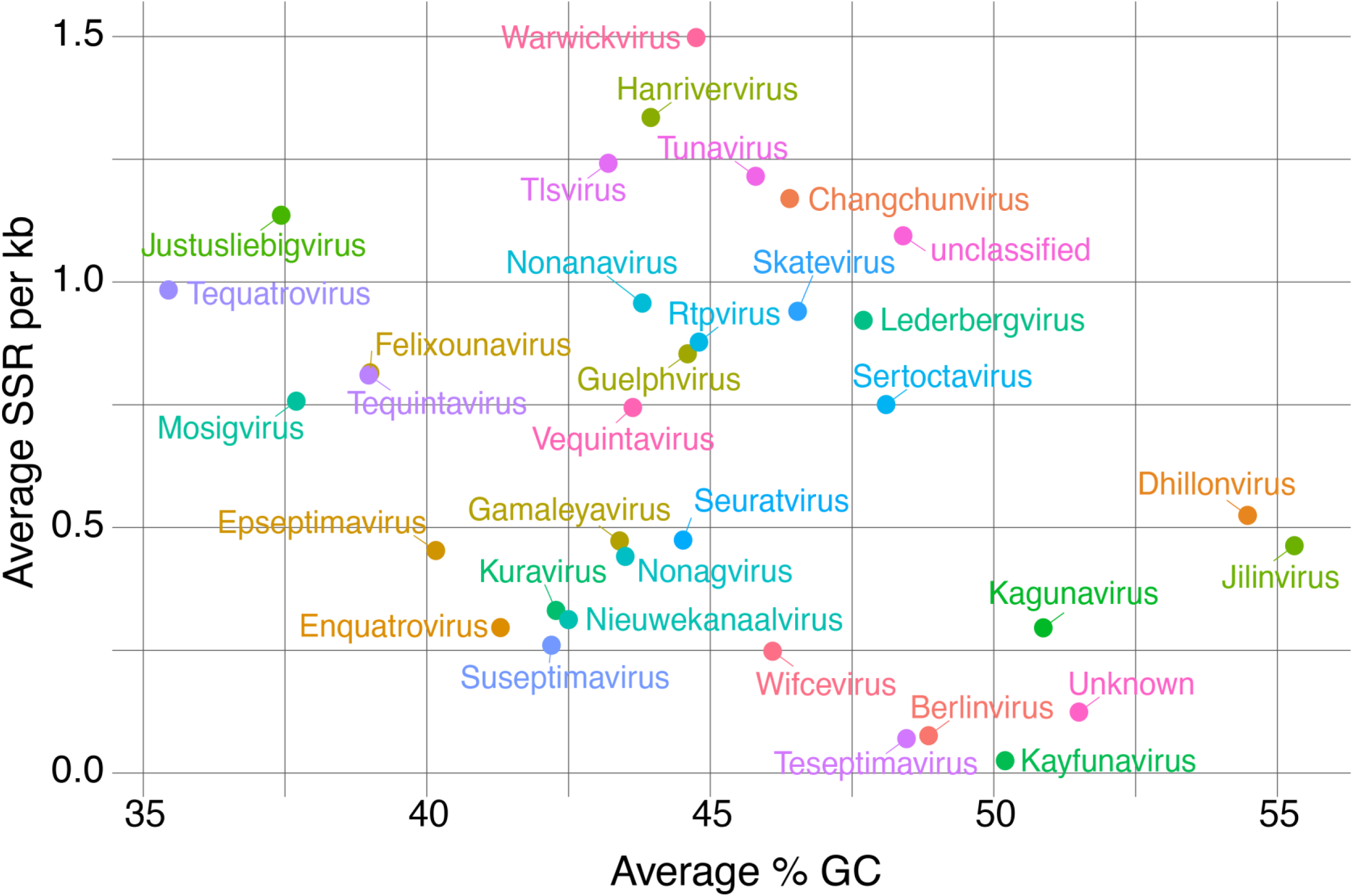
Simple sequence repeats ≥6 bases identified in other phages. Average putative CL per kb compared to average % GC of BASEL phage ICTV genus

**Extended Fig. 4.**
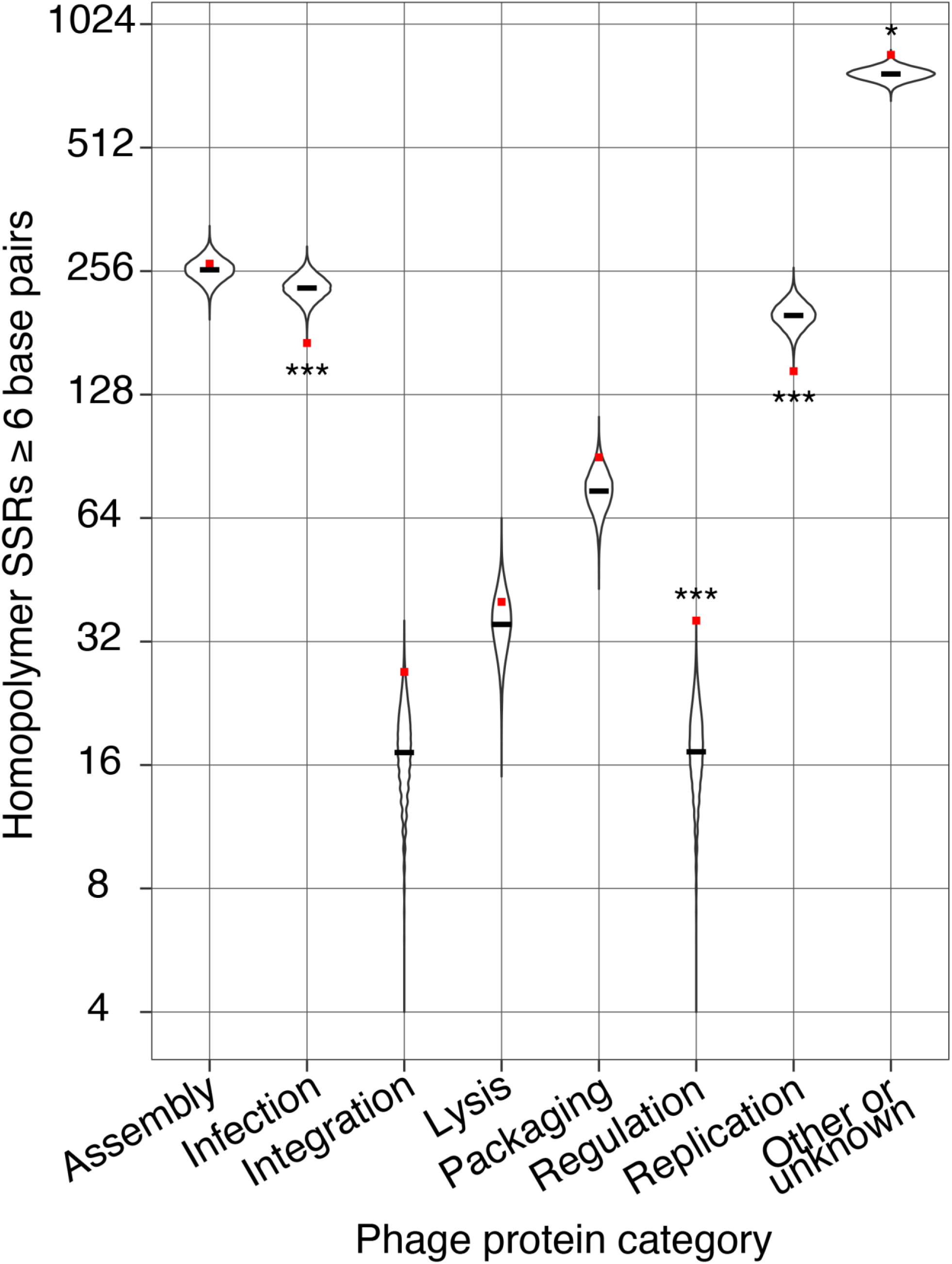
CL in BASEL phage collection. Homopolymer SSRs ≥6 base pairs identified in phage protein categories for 41 genetically distinct BASEL phage. Randomization test for each protein category was performed. Bonferroni adjusted p < 0.001 is indicated as ***, p = 0.014 is indicated as *

**Extended Fig. 5.**
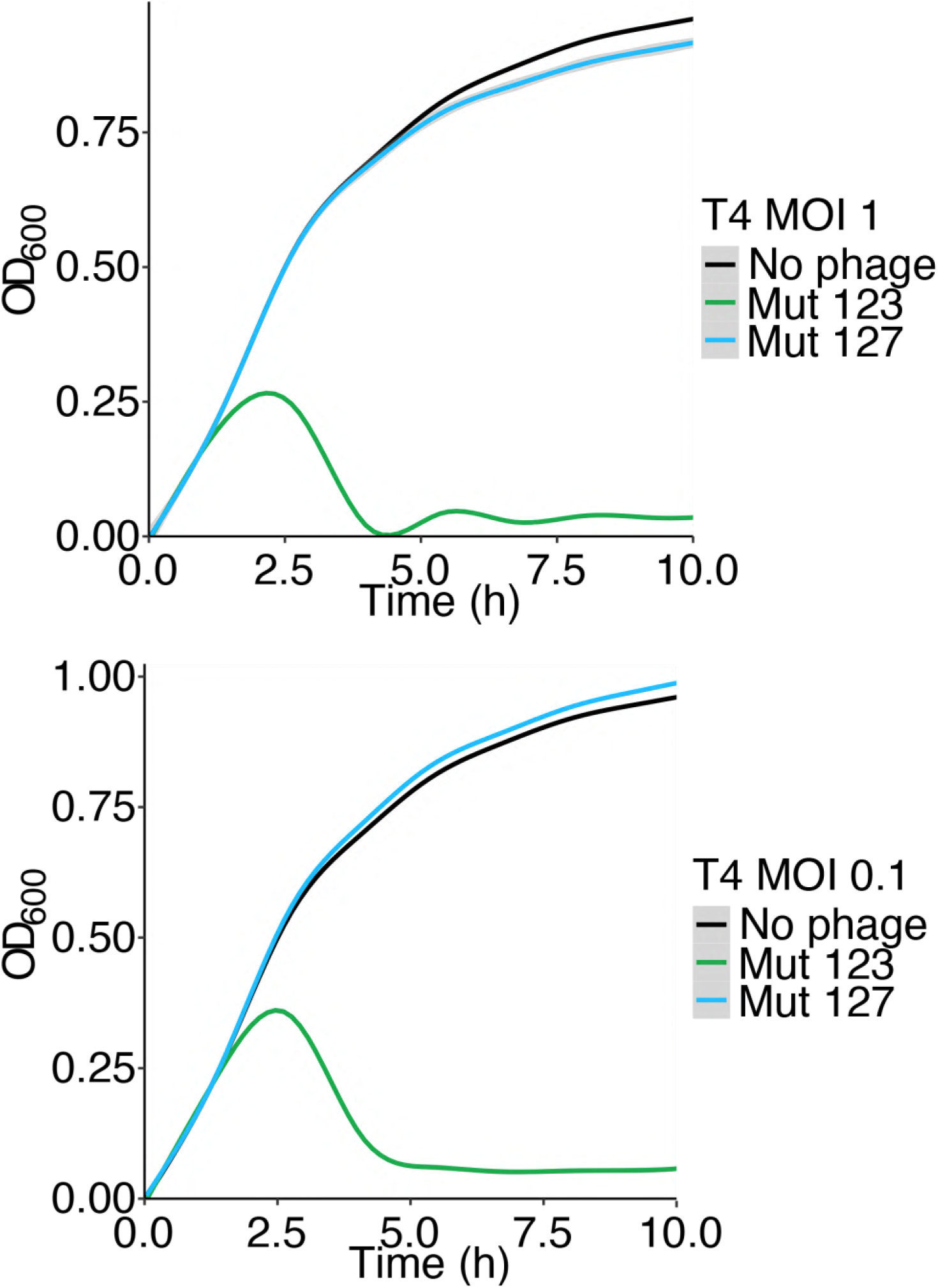
Evidence for CL reversion rate in T4 *rIIA*. MG1655(λ) infected with T4 Mut 123 or T4 Mut 127 at an MOI of 1 and 0.1 at time 0. The mean and standard error of 5 biological replicates each with 3 technical replicates are presented.

## References

1. Hendrix, R. W., Smith, M. C. M., Burns, R. N., Ford, M. E. & Hatfull, G. F. Evolutionary relationships among diverse bacteriophages and prophages: All the world’s a phage. Proc. Natl. Acad. Sci. 96, 2192–2197 (1999).

2. Wigington, C. H., Sonderegger, D., Brussaard, C. P. D., Buchan, A., Finke, J. F., Fuhrman, J. A., Lennon, J. T., Middelboe, M., Suttle, C. A., Stock, C., Wilson, W. H., Wommack, K. E., Wilhelm, S. W. & Weitz, J. S. Re-examination of the relationship between marine virus and microbial cell abundances. Nat. Microbiol. 1, 15024 (2016).

3. Moxon, E. R., Rainey, P. B., Nowak, M. A. & Lenski, R. E. Adaptive evolution of highly mutable loci in pathogenic bacteria. Curr. Biol. 4, 24–33 (1994).

4. Streisinger, G., Okada, Y., Emrich, J., Newton, J., Tsugita, A., Terzaghi, E. & Inouye, M. Frameshift mutations and the genetic code. Cold Spring Harb. Symp. Quant. Biol. 31, 77–84 (1966).

5. Levinson, G. & Gutman, G. A. Slipped-strand mispairing: a major mechanism for DNA sequence evolution. Mol. Biol. Evol. 4, 203–221 (1987).

6. Kroutil, L. C., Register, K., Bebenek, K. & Kunkel, T. A. Exonucleolytic proofreading during replication of repetitive DNA. Biochemistry 35, 1046–1053 (1996).

7. Attrill, E. L., Claydon, R., Łapińska, U., Recker, M., Meaden, S., Brown, A. T., Westra, E. R., Harding, S. V. & Pagliara, S. Individual bacteria in structured environments rely on phenotypic resistance to phage. PLoS Biol. 19, e3001406 (2021).

8. Kortright, K. E., Chan, B. K. & Turner, P. E. High-throughput discovery of phage receptors using transposon insertion sequencing of bacteria. Proc. Natl. Acad. Sci. 117, 18670–18679 (2020).

9. Mutalik, V. K., Adler, B. A., Rishi, H. S., Piya, D., Zhong, C., Koskella, B., Kutter, E. M., Calendar, R., Novichkov, P. S., Price, M. N., Deutschbauer, A. M. & Arkin, A. P. High-throughput mapping of the phage resistance landscape in *E. coli*. PLoS Biol. 18, e3000877 (2020).

10. Bernheim, A. & Sorek, R. The pan-immune system of bacteria: antiviral defence as a community resource. Nat Rev Microbiol 18, 113–119 (2020).

11. Millman, A., Melamed, S., Leavitt, A., Doron, S., Bernheim, A., Hör, J., Garb, J., Bechon, N., Brandis, A., Lopatina, A., Ofir, G., Hochhauser, D., Stokar-Avihail, A., Tal, N., Sharir, S., Voichek, M., Erez, Z., Ferrer, J. L. M., Dar, D., Kacen, A., Amitai, G. & Sorek, R. An expanded arsenal of immune systems that protect bacteria from phages. Cell Host Microbe 30, 1556–1569.e5 (2022).

12. Samson, J. E., Magadán, A. H., Sabri, M. & Moineau, S. Revenge of the phages: defeating bacterial defences. Nat. Rev. Microbiol. 11, 675–687 (2013).

13. Moxon, R., Bayliss, C. & Hood, D. Bacterial Contingency Loci: The role of simple sequence DNA repeats in bacterial adaptation. Genetics 40, 307–333 (2006).

14. Snowbarger, J., Koganti, P. & Spruck, C. Evolution of repetitive elements, their roles in homeostasis and human disease, and potential therapeutic applications. Biomolecules 14, 1250 (2024).

15. Vashakidze, R. P. & Prangishvili, D. A. Simple repetitive sequences in the genomes of archaebacteria. FEBS Lett. 216, 217–220 (1987).

16. Gomez, J. B. & Waters, C. M. A *Vibrio cholerae* Type IV restriction system targets glucosylated 5-hydroxymethylcytosine to protect against phage infection. J. Bacteriol. 206, e00143–24 (2024).

17. Vizzarro, G., Lemopoulos, A., Adams, D. W. & Blokesch, M. *Vibrio cholerae* pathogenicity island 2 encodes two distinct types of restriction systems. J. Bacteriol. 206, e00145–24 (2024).

18. Revel, H. R. & Luria, S. E. DNA-glucosylation in T-even phage: genetic determination and role in phage-host interaction 1. Annual Review of Genetics (1970).

19. Krüger, T., Wild, C. & Noyer-Weidner, M. McrB: a prokaryotic protein specifically recognizing DNA containing modified cytosine residues. EMBO J. 14, 2661–2669 (1995).

20. Sutherland, E., Coe, L. & Raleigh, E. A. McrBC: a multisubunit GTP-dependent restriction endonuclease. J. Mol. Biol. 225, 327–348 (1992).

21. Thomas, J. A., Orwenyo, J., Wang, L.-X. & Black, L. W. The odd “RB” phage— identification of arabinosylation as a new epigenetic modification of DNA in T4-like phage RB69. Viruses 10, 313 (2018).

22. Enikeeva, F. N., Severinov, K. V. & Gelfand, M. S. Restriction–modification systems and bacteriophage invasion: Who wins? J. Theor. Biol. 266, 550–559 (2010).

23. Georjon, H. & Bernheim, A. The highly diverse antiphage defence systems of bacteria. Nat. Rev. Microbiol. 1–15 (2023). doi:10.1038/s41579-023-00934-x

24. Boezen, D., Ali, G., Wang, M., Wang, X., Werf, W. van der, Vlak, J. M. & Zwart, M. P. Empirical estimates of the mutation rate for an alphabaculovirus. PLoS Genet. 18, e1009806 (2022).

25. Sanjuán, R., Nebot, M. R., Chirico, N., Mansky, L. M. & Belshaw, R. Viral mutation rates. J. Virol. 84, 9733–9748 (2010).

26. Stoler, N. & Nekrutenko, A. Sequencing error profiles of Illumina sequencing instruments. *NAR Genom. Bioinform.* 3, lqab019 (2021).

27. Schirmer, M., D’Amore, R., Ijaz, U. Z., Hall, N. & Quince, C. Illumina error profiles: resolving fine-scale variation in metagenomic sequencing data. BMC Bioinform. 17, 125 (2016).

28. Cowie, D. B., Avery, R. J. & Champe, S. P. DNA homology among the T-even bacteriophages. Virology 45, 30–37 (1971).

29. Luria, S. E., Delbrück, M. & Anderson, T. F. Electron microscope studies of bacterial viruses. J. Bacteriol. 46, 57–77 (1943).

30. Kim, J.-S. & Davidson, N. Electron microscope heteroduplex study of sequence relations of T2, T4, and T6 bacteriophage DNAs. Virology 57, 93–111 (1974).

31. Molineux, I. J. The T7 Group. The Bacteriophages: Second Edition 0 (2005). doi:10.1093/oso/9780195148503.003.0020

32. Doron, S., Melamed, S., Ofir, G., Leavitt, A., Lopatina, A., Keren, M., Amitai, G. & Sorek, R. Systematic discovery of antiphage defense systems in the microbial pangenome. Science 359, (2018).

33. Seed, K. D., Bodi, K. L., Kropinski, A. M., Ackermann, H.-W., Calderwood, S. B., Qadri, F. & Camilli, A. Evidence of a dominant lineage of *Vibrio cholerae*-specific lytic bacteriophages shed by Cholera patients over a 10-year period in Dhaka, Bangladesh. Mbio 2, e00334–10 (2011).

34. Hays, S. G. & Seed, K. D. Dominant Vibrio cholerae phage exhibits lysis inhibition sensitive to disruption by a defensive phage satellite. eLife 9, e53200 (2020).

35. Madi, N., Cato, E. T., Sayeed, M. A., Creasy-Marrazzo, A., Cuénod, A., Islam, K., Khabir, M. I. U., Bhuiyan, M. T. R., Begum, Y. A., Freeman, E., Vustepalli, A., Brinkley, L., Kamat, M., Bailey, L. S., Basso, K. B., Qadri, F., Khan, A. I., Shapiro, B. J. & Nelson, E. J. Phage predation, disease severity, and pathogen genetic diversity in cholera patients. Science 384, (2024).

36. Wang, R. H., Yang, S., Liu, Z., Zhang, Y., Wang, X., Xu, Z., Wang, J. & Li, S. C. PhageScope: a well-annotated bacteriophage database with automatic analyses and visualizations. Nucleic Acids Res. 52, D756–D761 (2023).

37. Humolli, D., Piel, D., Maffei, E., Heyer, Y., Agustoni, E., Shaidullina, A., Willi, L., Imwinkelried, P., Estermann, F., Cuénod, A., Buser, D. P., Alampi, C., Chami, M., Egli, A., Hiller, S., Dunne, M. & Harms, A. Completing the BASEL phage collection to unlock hidden diversity for systematic exploration of phage–host interactions. PLOS Biol. 23, e3003063 (2025).

38. Crick, F. H. C., Barnett, L., Brenner, S. & Watts-Tobin, R. J. General nature of the genetic code for proteins. Nature 192, 1227–1232 (1961).

39. Fisher, K. M. & Bernstein, H. The additivity of intervals in the rIIA cistron of phage T4D. Genetics 52, 1127–1136 (1965).

40. Benzer, S. Fine structure of a genetic region in bacteriophage. Proc. Natl. Acad. Sci. 41, 344– 354 (1955).

41. Benzer, S. On the topography of the genetic fine structure. Proc. Natl. Acad. Sci. 47, 403– 415 (1961).

42. Edgar, R. S., Feynman, R. P., Klein, S., Lielausis, I. & Steinberg, C. M. Mapping experiments with r mutants of bacteriophage T4D. Genetics 47, 179–186 (1962).

43. Shinedling, S., Singer, B. S., Gayle, M., Pribnow, D., Jarvis, E., Edgar, B. & Gold, L. Sequences and studies of bacteriophage T4 rII mutants. J. Mol. Biol. 195, 471–480 (1987).

44. Wong, S., Alattas, H. & Slavcev, R. A. A snapshot of the λ T4rII exclusion (Rex) phenotype in *Escherichia coli*. Curr. Genet. 67, 739–745 (2021).

45. Shinedling, S., Parma, D. & Gold, L. Wild-type bacteriophage T4 is restricted by the lambda rex genes. J. Virol. 61, 3790–3794 (1987).

46. Paddison, P., Abedon, S. T., Dressman, H. K., Gailbreath, K., Tracy, J., Mosser, E., Neitzel, J., Guttman, B. & Kutter, E. The roles of the bacteriophage T4 r genes in lysis inhibition and fine-structure genetics: A new perspective. Genetics 148, 1539–1550 (1998).

47. Sørensen, M. C. H., Vitt, A., Neve, H., Soverini, M., Ahern, S. J., Klumpp, J. & Brøndsted, L. *Campylobacter* phages use hypermutable polyG tracts to create phenotypic diversity and evade bacterial resistance. Cell Rep. 35, 109214 (2021).

48. Liu, M., Gingery, M., Doulatov, S. R., Liu, Y., Hodes, A., Baker, S., Davis, P., Simmonds, M., Churcher, C., Mungall, K., Quail, M. A., Preston, A., Harvill, E. T., Maskell, D. J., Eiserling, F. A., Parkhill, J. & Miller, J. F. Genomic and genetic analysis of *Bordetella* bacteriophages encoding reverse transcriptase-mediated tropism-switching cassettes. J. Bacteriol. 186, 1503– 1517 (2004).

49. Ceyssens, P., Miroshnikov, K., Mattheus, W., Krylov, V., Robben, J., Noben, J., Vanderschraeghe, S., Sykilinda, N., Kropinski, A. M., Volckaert, G., Mesyanzhinov, V. & Lavigne, R. Comparative analysis of the widespread and conserved PB1-like viruses infecting *Pseudomonas aeruginosa*. Environ. Microbiol. 11, 2874–2883 (2009).

50. Yen, M., Cairns, L. S. & Camilli, A. A cocktail of three virulent bacteriophages prevents *Vibrio cholerae* infection in animal models. Nat. Commun. 8, 14187 (2017).

51. Deatherage, D. E. & Barrick, J. E. Engineering and analyzing multicellular systems, methods and protocols. Methods Mol. Biol. 1151, 165–188 (2014).

52. Barrick, J. E. & Zubbu, I. Brefito: Snakemake pipelines for analyzing resequenced bacterial genomes. Brefito: Snakemake pipelines for analyzing resequenced bacterial genomes. (2025). at <https://github.com/barricklab/brefito>

53. Cock, P. J. A., Antao, T., Chang, J. T., Chapman, B. A., Cox, C. J., Dalke, A., Friedberg, I., Hamelryck, T., Kauff, F., Wilczynski, B. & Hoon, M. J. L. de. Biopython: freely available Python tools for computational molecular biology and bioinformatics. Bioinformatics 25, 1422– 1423 (2009).

54. Jain, C., Rodriguez-R, L. M., Phillippy, A. M., Konstantinidis, K. T. & Aluru, S. High throughput ANI analysis of 90K prokaryotic genomes reveals clear species boundaries. Nat. Commun. 9, 5114 (2018).

55. Camacho, C., Coulouris, G., Avagyan, V., Ma, N., Papadopoulos, J., Bealer, K. & Madden, T. L. BLAST+: architecture and applications. BMC Bioinform. 10, 421 (2009).

